# FCS in closed systems and application for membrane nanotubes

**DOI:** 10.1101/134742

**Authors:** Yanfei Jiang, Artem Melnykov, Elliot E. Elson

## Abstract

In the present study, we developed the fluorescence correlation spectroscopy theory for closed systems with either periodic or reflective boundaries. The illumination could be any arbitrary function. We tested our theory with simulated data of both boundary conditions. We also tested the theory with experimental data of membrane nanotubes, whose circular direction is a closed system. The result shows that the correlation function for nanotubes falls between 1D and 2D diffusion model. The fitting with our model gives an accurate recovery of the diffusion time and nanotube radius. We also give some examples of single molecule experiments for which our theory can be potentially useful.

## 1. Introduction

Lipid membranes can form different structures in cells. One of structures is the recently discovered tunneling nanotubes (TNTs). Transportation through the tunneling nanotubes is believed to be a new cell-cell communication mechanism (Davis and Sowinski 2008; Gerdes and Carvalho 2008). These nanotubes are also found to be important in HIV virus spreading among cells. The diameter of TNTs is between 50 and 400 nm, while the length can reach from several to hundreds of micron(Davis and Sowinski 2008). They have been found in a variety of cell types, such as epithelial cells (Lehmann, Sherer et al. 2005; Sherer, Lehmann et al. 2007), immune cells (Onfelt, Nedvetzki et al. 2004; Watkins and Salter 2005; Onfelt, Nedvetzki et al. 2006; Sowinski, Jolly et al. 2008), and neural cells (Rustom, Saffrich et al. 2004; Wang, Cui et al. 2011). This indicates that transportation through the TNTs is a general physical communication mechanism for cells separated with long-distances.

A great amount of interest in TNTs has focused on the traffic of proteins and organelles between cells, either inside of the TNT tunnels or in the TNT membranes (Davis and Sowinski 2008; Gerdes and Carvalho 2008; Suhail, Kshitiz et al. 2013). There are still many questions remaining about the trafficking mechanism and TNT formation. The formation of some of the special membrane structures, like the virus bud, involves not only the clustering of the relevant proteins, but also certain kinds of lipids to form lipid rafts (Nguyen and Hildreth 2000; Simons and Vaz 2004; Rajendran and Simons 2005). In the present study, we are also interested in whether the membrane properties are the same between the cell body’s membrane and the TNT membrane. Fluorescence correlation spectroscopy has been widely used to study the diffusivity of the molecules in the membrane as we have discussed in the previous chapters. It is a potentially powerful tool to study the trafficking in the TNT membrane and the membrane properties of TNTs.

A straightforward assumption with FCS measurements on TNTs is to consider the TNTs as a one-dimensional model. However, since the diameter of the TNTs is comparable to the size of the observation area of FCS, it is necessary to investigate the dependence of the diffusion behavior on the TNT diameter. Unlike traditional FCS which usually deals with open systems, the tube structure can be regarded as open system only in the direction along the axis. For the direction around the axis, it is a closed system with periodic boundary conditions. In the present study, we first derived the general expressions for FCS in the closed systems and then applied them to the FCS measurements in the tubular membranes. We found the correlation magnitude between the closed systems and the corresponding open systems is differed by a value which equals to the inverse of the number of molecules in the system. We also found that one-dimensional simplification is only good for tubes whose radius is less than 80 nm with the laser waist being 250 nm. Our theoretical calculation also suggests that the apparent correlation time decreases with the increasing tube radius, which is anti-intuitive. The result is supported by a monte-carlo simulation.

## 2. Material and Methods

### 2.1 Lipids and GUV preparation

All the lipids were purchased from Avanti Polar Lipids. DiI-C20 was purchased from Molecular Probes. Lipids were stocked at 10 mM in chloroform and the DiI-C20 was in ethanol at 0.01 mM. GUVs were made by electroformation (Morales-Penningston, Wu et al. 2010; Walde, Cosentino et al. 2010). DiD-C20 stock solutions were first mixed with DLPC solution and deposited onto two platinum wires by dragging the lipid solution along the wires back and forth until the chloroform and ethanol dried. Then the two wires were put into a home-made Teflon cylinder chamber filled with 300 mM sucrose solution. The chamber cap has two holes to hold the wires and separate them at a distance of 2 mm. The chamber was next put at room temperature or on a heating stage, which keeps the temperature at 70 ^°^C. The wires were then connected to a function generator giving a 10 Hz/1.5 volt square wave for 1-2 hour.

### 2.2 Nanotube preparation

The Lab-tek 8 well chamber was used for nanotube experiments. The chamber glass was first treated with poly-lysine (0.1% w/v) for 1 hour. 200 *μl* water was first added to the chamber and the chamber with water was then heated to 60 degree before the GUVs in the Teflon chamber were rapidly transferred. After transferring, the chamber was then put in the 60 degree oven for 5 minute to allow the GUVs sediment to the bottom. Then the chamber was taken out and shaken laterally by hand. Because the bottom of some of the GUVs will attached to the glass surface, nanotubes form when these GUVs move with chamber shaking.

### 2.3 Fluorescence microscopy and FCS measurements

Fluorescence microscopy and FCS measurements were performed on a LSM 510 ConfoCor 2 CombinationSystem from Carl Zeiss, Germany. The system is based on a Zeiss Axiovert 200M inverted microscope. A C-Apochromat 40X objective (numerical aperture 1.2) was used. GUVs and nanotubes were first found in the imaging mode and then moved using the microscope stage to the position defined for FCS measurements. For the measurements on GUVs, a z-scan was first performed to find the apex of the vesicle. For the measurements on nanotubes, both z-scan and x-scan are needed to position the nanotube in the center of the exitation laser. 5-20 FCS measurements were performed for each GUV. The FCS curves were fitted using home-made Matlab programs.

## 3. Theory

### 3.1 FCS for the closed systems

The (temporal) autocorrelation function, which is the correlation of a time series with itself shifted by time *τ*, is calculated as following:

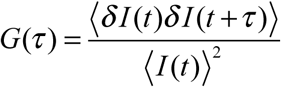

〈*I*(*t*)^2^ is the square of the mean fluorescence, considered as a normalization factor in the calculation of the autocorrelation function. In the theory of FCS, it can be expressed as following.

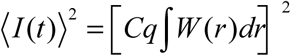

*C* the concentration of the fluorescence particles and *q* is the brightness per molecule. *W*(*r*) is the detectable emission intensity distribution (spatial detectivity function). It can be noticed that 〈*I*(*t*)^2^ is independent of the lag time τ.

In the following context, we use lower case *g*(*τ*) to represent the unnormalized autocorrelation function. The standard unnormalized autocorrelation function for open systems is calculated as follows.

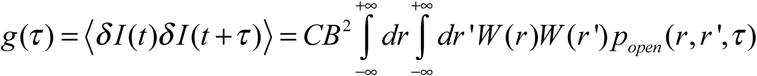

C is the concentration of molecules and B is the brightness per molecule. *p*_*open*_ (*r*, *r* ',*τ*) is called the number density autocorrelation, which is the probability density for a single molecule that started a random walk at time 0 at the point *r* to be at *r’* at a lag time, and it can be expressed as (Schwille, Korlach et al. 1999).

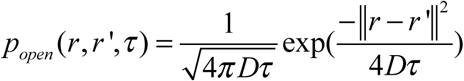

We can see that *p*_*open*_ (*r*, *r* ',*τ*) approaches 0 as the lag time goes to infinity. This is because the probability of one molecule returning the same position is zero at the infinite time when the system is open. Therefore, *G*_*open*_ (∞) = 0. At the other end, when *r=r’*, and τ= 0, *p*_*open*_ (*r*, *r'*,*τ*) approaches the Dirac delta function. Therefore, *g*(0) = *Cq*^2^ ∫*W*(*r*)^2^*dr*, and 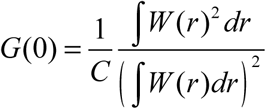. The term 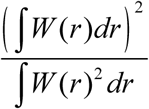 is often called effective volume, *V*_*effective*_.

**Figure 2.**
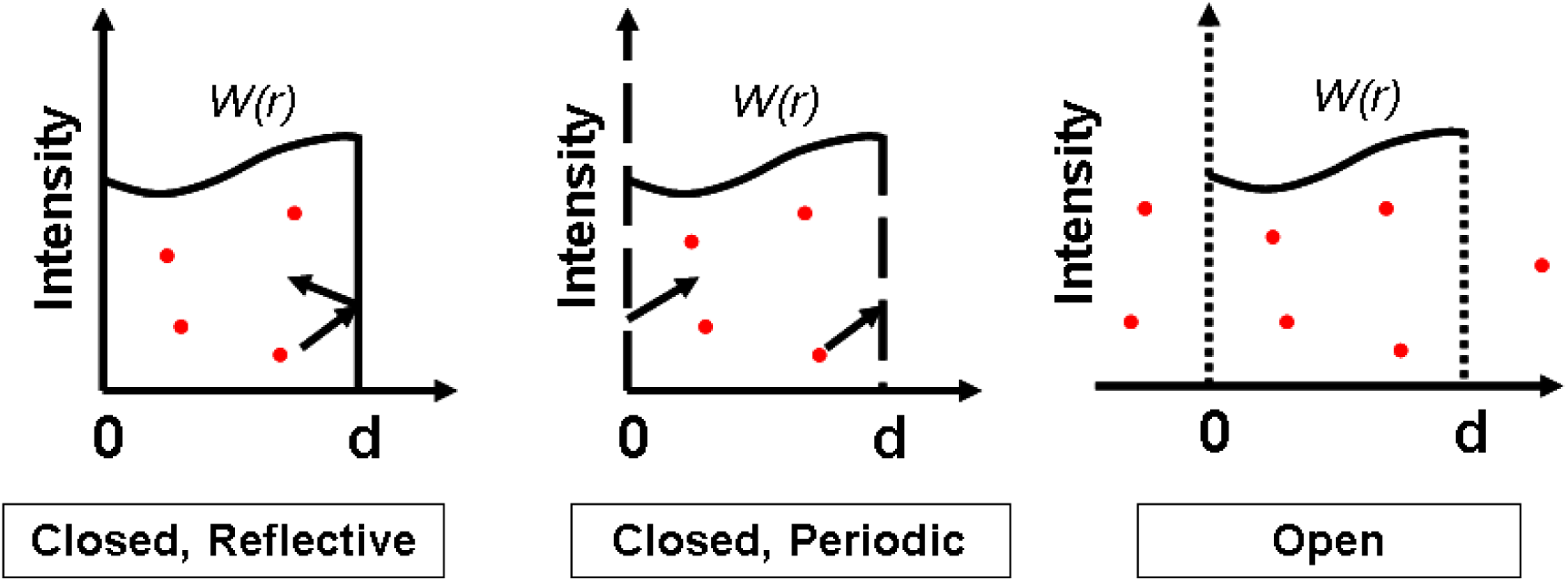
Illustration of the closed systems and open systems. The boundary condition can be either reflective and periodic for the closed systems. The curve is the illumination profile *W*(*r*).

For closed systems, molecules can not diffuse to infinity, therefore,

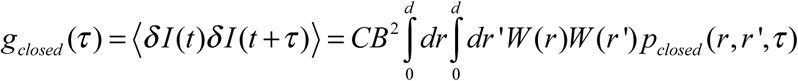

There are two different boundary conditions, periodic and reflective. For reflective boundaries, the molecules are reflected back into the system once they diffuse into the system boundary “wall”. Periodic boundary conditions are used to account for diffusion that arrives at its starting point after traversing the whole system. This usually happens in a ring structure. For both of the two conditions, the number density autocorrelation can be derived using the superposition method which involves folding the unconfined number density autocorrelation, *p*_*open*_ (*r*, *r'*,*τ*), along the system walls. The confined probability for the closed systems is then obtained by summing up all the parts derived from the reflections or reenterings.

Therefore,

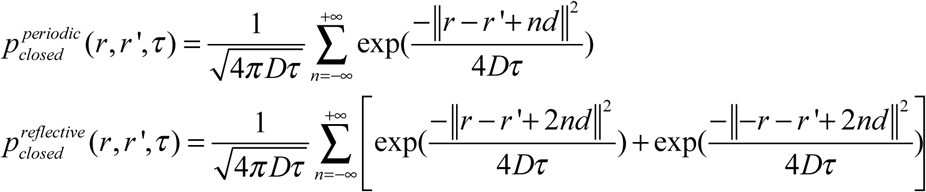

*d* is the size of the closed system.

### 3.2 Amplitude of the correlation

For equation and, both these two probability density distributions approach the Dirac delta function when *r=r’*, and *τ* = 0. On the other hand, when *τ* → ∞,

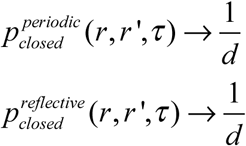

This means the probabilities of a molecule starting from one position and ending at any positions in the system are identical and constant. This constant probability is just the inverse of the size of the system.

For closed systems, when *r=r’*, and τ = 0, *p*_*open*_ (*r*, *r'*,*τ*) also approaches the Dirac delta function as the open systems do. Therefore, *g*(0) = *Cq*^2^ ∫*W*(*r*)^2^ *dr*, and 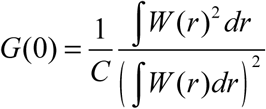 However, at the infinite lag time, because the 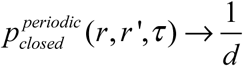,

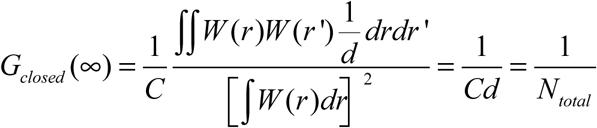

*N*_*total*_ is the total number of particles in the system. It can only take positive integer number values for a closed system, while the average number of particles in the excitation area for the open systems can be any positive number. Because *G*(0) is the same for both closed and open systems, the magnitude of the correlation function (*G*(0) – *G*(∞)) is different by a factor of 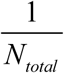 when the average number of the particles in the open systems is the same as the total number of particles in the corresponding closed systems. The smaller magnitude for the closed systems can be attributed to the fact that the fluorescence fluctuation is only caused by the unevenness of the*W*(*r*). When *W*(*r*) is constant, there will not be any fluctuation and the correlation goes to zero for the closed systems. However, this is not the case for the open systems because the number of particles still fluctuates around the average, resulting in fluorescence fluctuation and positive correlation.

For a simple one-component system, the fluorescence fluctuation is from three major sources: (1) dye concentration fluctuation; (2) nonuniform illumination or unevenness of illumination; (3) shot noise. Since shot noise is not correlated, we will next only discuss the first two kinds of sources. On one hand, for closed systems, the total number of molecules is constant, so the fluctuation is only caused by the unevenness of illumination. As we have shown above, the zero time magnitude of the correlation function is different by a factor of 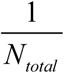 for the closed and open systems. On the other hand, for an open system that has uniform illumination, the fluctuation is only induced by the concentration fluctuation. It can be easily shown that the zero time magnitude of the correlation function for such an open system is 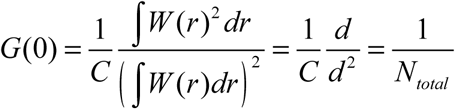, which is exactly the difference between the closed system and the open system. Therefore, the magnitude of the correlation function *G*(0) – *G*(∞) for any open system is the sum of the magnitudes of the correlation function for the corresponding closed system and for the corresponding uniform illumination open system. This provides a way to quantitatively elucidate the contributions by the two sources for the fluorescence fluctuation. Briefly, the contribution by the unevenness of illumination is 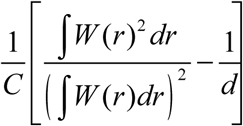 and the contribution by dye concentration fluctuation is 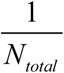.

### 3.3 Evaluation the correlation function

#### 3.3.1 In real space

Instead of calculating the correlation function by folding up the number density autocorrelation function, we can also keep this function extended but also extend the illumination to infinity, as depicted in Figure 3. By doing this, the complicated summing up in equations and can be avoided. The correlation function can be expressed by

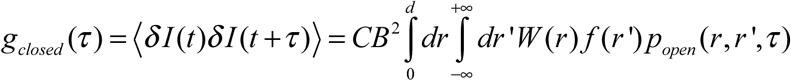

*f*(*r*) is the extended version of *W*(*r*). For periodic boundaries, *f*(*r*) is the repeat of *W*(*r*) and the period of *f*(*r*) is *d*. For reflective boundaries, *f*(*r*) is the repeat of *W*(*r*) and its image. Therefore, the period of *f*(*r*) is 2*d.*

**Figure 3.**
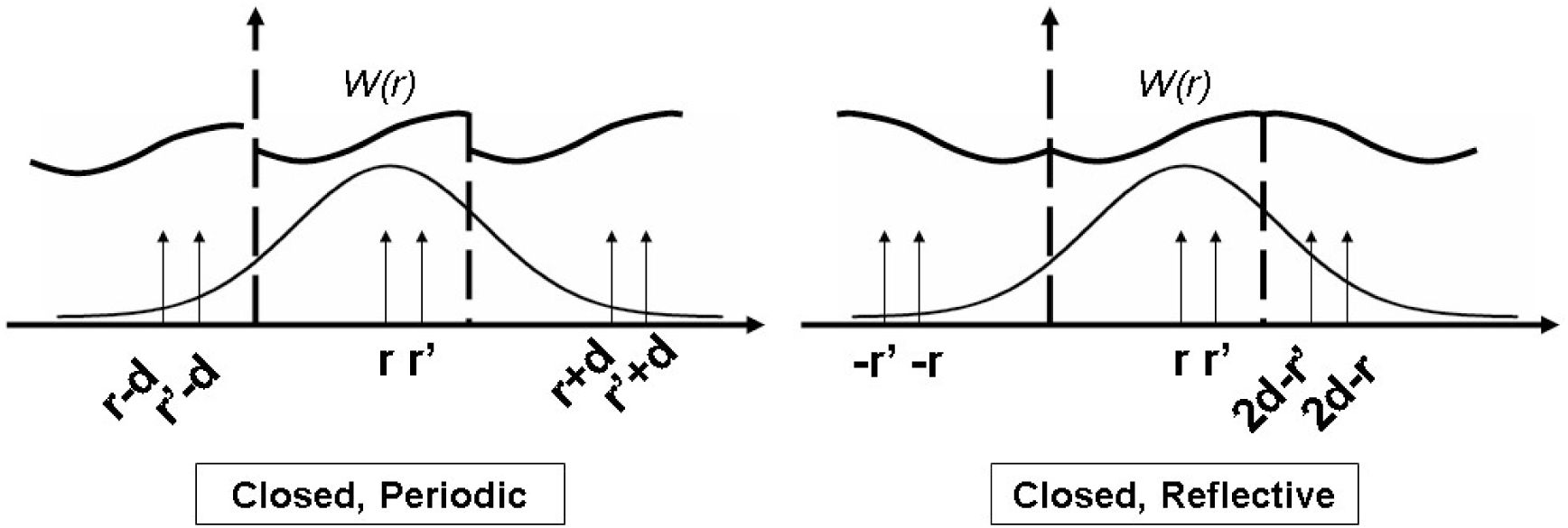
Unfold the closed system along the wall. The Gaussian curve is the number density autocorrelation.

#### 3.3.2 In Fourier space

The double integration in equation still takes a relatively long time to evaluate and is not suitable for curve fitting. In order to accelerate the calculation, Fourier transform method can be used, by which the double integral can be changed into a single integral. For open systems, (Elson and Webb 1975; Koppel, Axelrod et al. 1976)

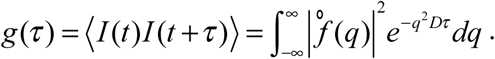

°*f*(*q*) is the Fourier transform of the beam shape, 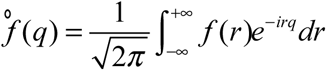.

Sanguigno L et al (Sanguigno, De Santo et al. 2010) have derived the expression for a confined box system using the Fourier transform method. Here, we generalize the method for both reflective and periodic boundary conditions and provide a basic framework for any beam profile (see Appendix for derivations):

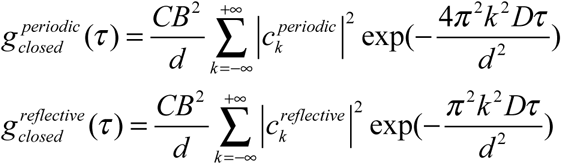

The factors 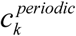 and 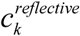 are given by:

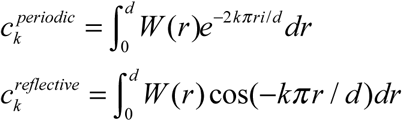

Therefore, the normalized correlation functions are:

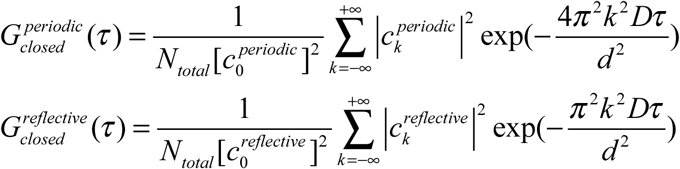

The factors 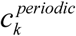 and 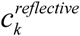are independent of the lag time and diffusion rate, so they can be pre-calculated to accelerate the function evaluation. Unlike the Gaussian laser, most *W*(*r*) s do not have explicit expressions for the correlation functions; evaluation with the Fourier transform method is still much faster than the double integration in equation.

As discussed above, the total number of molecules for a closed system only takes integer numbers, so whether the total number of molecules is close to an integer indicates whether the selected *W*(*r*) is accurate. One common reason that makes the *W*(*r*) inaccurate is the background fluorescence. This can be solved by adding a background term to the *W*(*r*). If the background is uniform, it only changes the value of 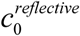 or 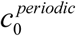. Therefore, instead of recalculating all the 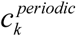 and 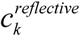 for different *W*(*r*) s, a single parameter can be added to 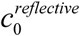 or 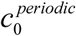 for evaluation.

### 3.4 FCS in rings and tubes

For tubular membranes placed in the center of the focused laser and parallel to the surface (xy plane) as shown in Figure 1, the system can be considered as open in the y direction and closed in the xz plane. In the y-direction, the correlation function can be shown as:

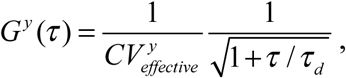

where 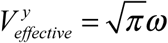 and 2*Dτ*_*d*_ = *w*^2^

**Figure 1.**
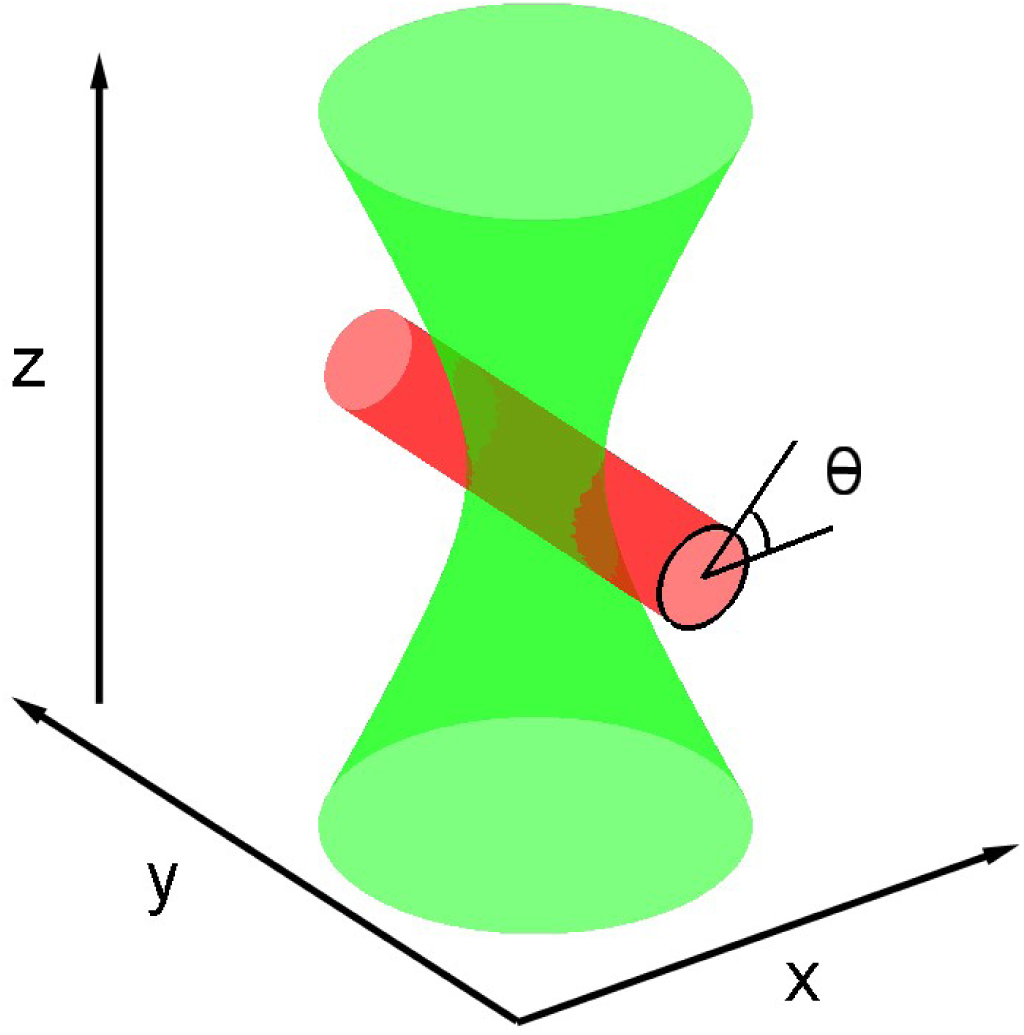
FCS measurement in the nanotube membrane. The molecules in the membrane diffuse in two directions. One is the open direction in y and the other is the closed direction around the tube.

We will focus on the closed direction in the following paragraphs. In the xz plane, the molecules diffuse in a ring, meaning this system has a periodic boundary condition. The size of the system is *d* = 2*πR*, where *R* is the radius of the ring/tube. Since we are interested in nanotubes, whose diameters are much smaller than the size of beam in the z direction, we consider the excitation in z to be constant. Therefore,

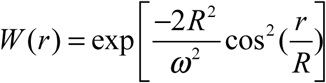

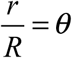, as shown in Figure 1.

With the size of the system and the laser profile, the correlation function can be obtained for different diffusion coefficients using the Fourier transform method described above. Particularly, we want to discuss the zero time magnitude of the correlation which is independent of the diffusion rate. Although the full correlation function does not have an explicit expression, the zero time magnitude does.

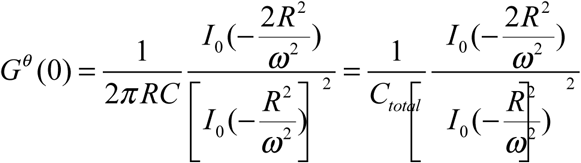

*I*_0_ is the modified Bessel function of first kind. It can also be expressed as:

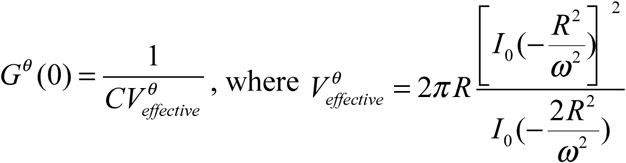

For the tube structure, the correlation function is the product of the correlation functions in both directions.

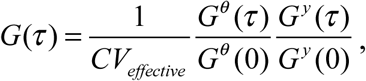

Here, 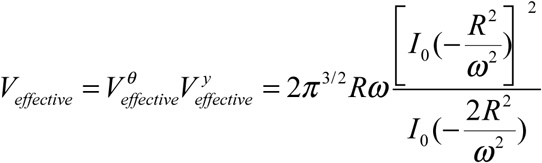.

## 4. Results

### 4.1 FCS for rings

Figure 4 shows the correlation function for diffusion in a ring. Both theoretical and simulated results are shown. There’s one thing that should be noticed: The ACF decay to zero has to be imposed by subtracting the *G*(∞). Sanguigno L argued that in the open systems, any kind of fluctuation, considered as an initial condition, approaches zero at infinite time. However, in closed systems, the ACF decay to zero has to be imposed by subtracting the *G*(∞) since it is not always true for all possible initial conditions (Sanguigno, De Santo et al. 2010). We also provide an alternative explanation in the section of PCH and FIDA for closed system.

**Figure 4.**
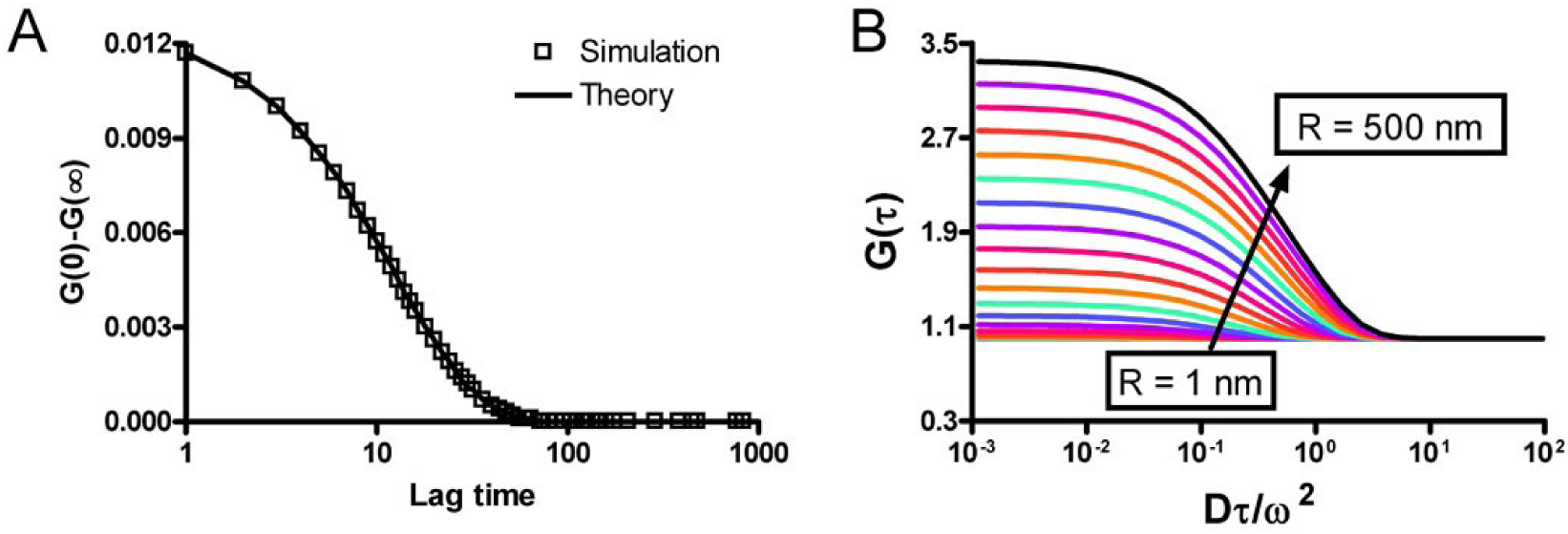
Comparison between the theoretical and the simulated correlation functions for the FCS in a ring. The ring radius is 100 nm and the laser waist is 250 nm. The center of the ring is place in the middle of the laser.

We can see that the theory predicts the correlation function very accurately. In the simulation, there is only one molecule in the ring. For an open system whose average number of molecules is one, the magnitude of the correlation should be the 1 / *V*_*effective*_, which is 1.012 according to equation 7. But as shown in Figure 4, the magnitude for the closed system is 0.012, which is much smaller than the open system. It means in the FCS measurement, the contribution by the number fluctuation is much large than the contribution by the unevenness of the illumination.

### 4.2 FCS for tubes

We calculated the theoretical correlation functions (Figure 5, left) for different ring radii with the identical beam size, *ω* = 250 nm. As shown in Figure 5, the correlation function (normalized by the G(0)) is close to 1 for a small radius, which means the system fluctuates less. Secondly, the lag time for the correlation to decrease to half magnitude becomes longer.

**Figure 5.**
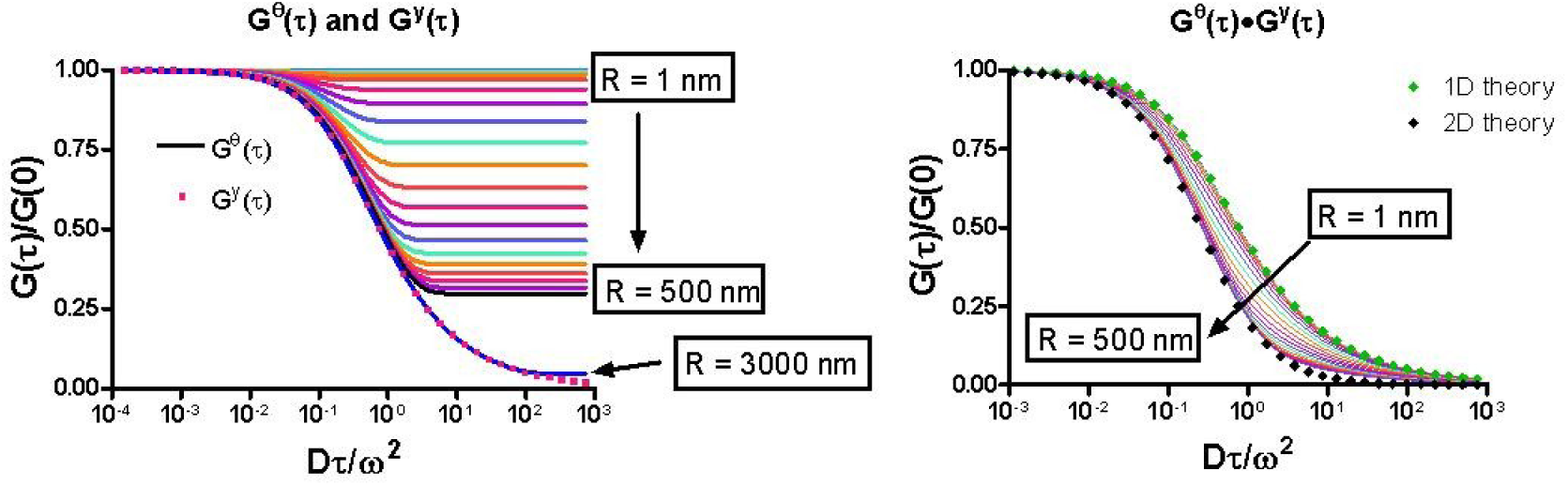
Theoretical correlation functions for nanotubes. (left) the correlation function for different ring sizes (lines), compared to the correlation function in the y direction which is open direction (pink dot). (right) the product of the correlation functions in the two directions (lines). For comparison, the correlation function for simple 1D and 2D systems are also shown in green and black dots.

This is caused by the fact that it takes longer for molecules to diffuse around bigger ring sizes. Overall, the decay time in the ring is smaller than the decay time in the open dimension. Thirdly, because the curvature of the ring becomes more and more flat with increasing radius, the diffusion in the ring will approach 2D diffusion with large ring radius.

This is supported by our results. As shown in Figure 5 left, when the R = 3000 nm, the *G*^*θ*^ (*τ*) is almost identical to the *G* ^*y*^ (*τ*).

The right panel of Figure 5 shows the result for the real tube system, whose correlation function is the product of the correlation functions of the closed ring and the open dimension. Our simulation shows that the theory predicts the correlation function very well for the tube membrane system (Figure 6). In order to understand the system, we are interested in two extremes. At one extreme point, when the tube radius is zero, *G*^*θ*^ (*τ*) will constantly equal to 1, leading to *G*(*τ*) = *G* ^*y*^ (*τ*). This means the system is just one dimensional. At the other extreme point, when the radius is infinitely large, *G*^*θ*^ (*τ*) will equal to *G* ^*y*^ (*τ*) and therefore, the system will be a two dimensional system. Our theoretical calculation shows this gradual transition from one-dimension to two-dimensions as the tube radius increases.

**Figure 6.**
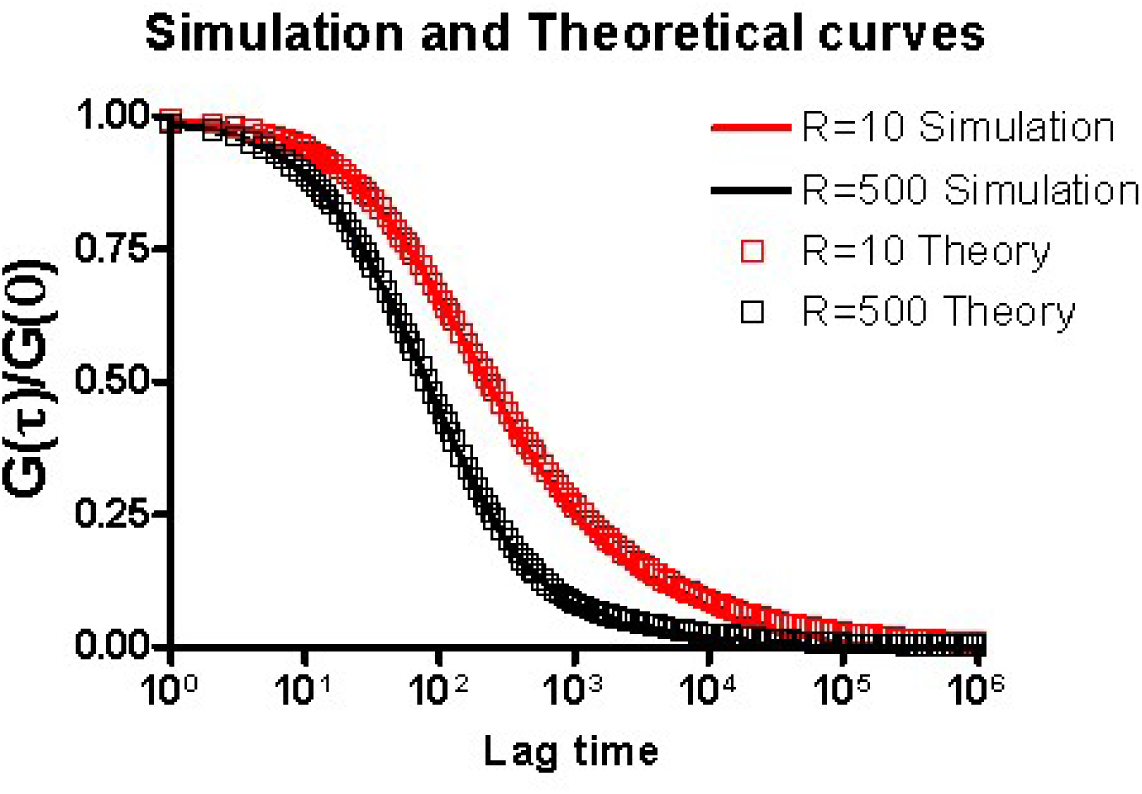
Comparison between the theoretical and the simulated correlation functions for the FCS in a nanotube. The laser waist is 250 nm. The tube radii are 10 and 500 nm.

One thing to notice is that the apparent correlation time (roughly the lag time that the correlation decreases to half of magnitude) changes in opposite directions for the ring and the tube. This means for a multi-dimensional system, the change of the correlation time in one of the dimensions does not necessarily causes the same change in the overall correlation time. In the current case, the decrease of the overall correlation time is due to the switch from one-dimensional diffusion to two-dimensional diffusion.

For the nanotube membranes, we would like to know whether we can use the simplified one-dimension model to fit the experimental data. In order to answer this question, we used the one dimension model to fit the theoretical correlation function for the nanotubes. The fitted correlation time and *χ* ^2^ are plotted in Figure 7. As we expected, the fitted correlation time gradually decreases with the increasing radius and the *χ* ^2^ increases. It is also obvious the confidence interval of the fitted correlation time gets bigger. These results all suggest the unsuitability of the one-dimension model for large radius. For the *χ* ^2^, the main turning point is at the 200 nm. For tubes larger than 200 nm, the *χ* ^2^ was obviously large. However, the fitted correlation time is only about 0.6 fold of the real value for 200 nm tubes. The turning point for the fitted correlation time is smaller, which is around 80 nm. So we conclude that, for nanotubes whose radius is less than 80 nm, it is sufficient to simplify the system and use the one dimensional model. However, for nanotubes larger than 80 nm, it is recommended to use the more complete model which takes the fluctuation in the closed ring into account.

**Figure 7.**
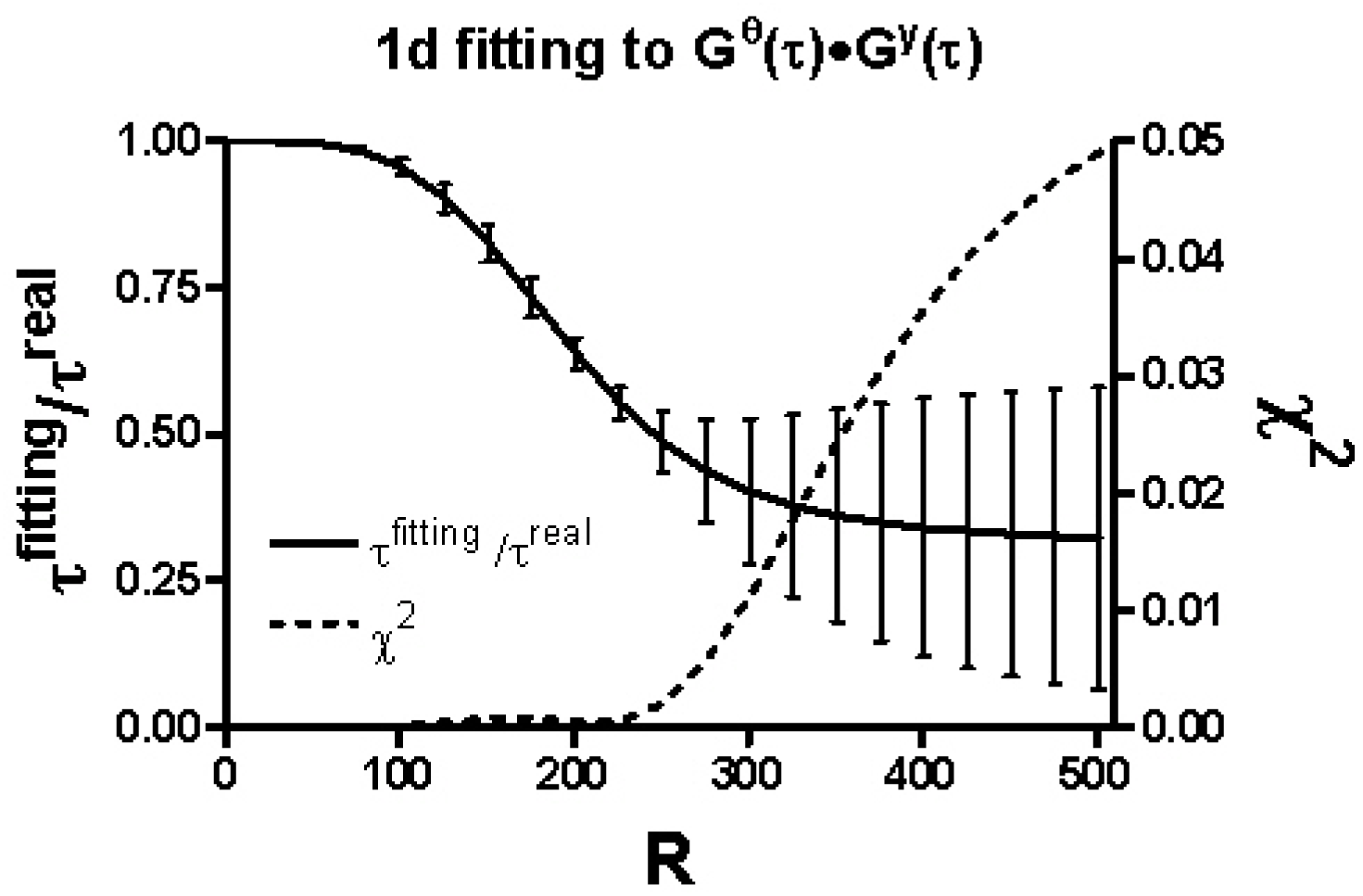
Fitting results to the theoretical correlation functions for the nanotube using 1D model.

### 4.3 Experimental test of the FCS for nanotubes

To test our theory, we made some lipid membrane nanotubes and conducted FCS experiments on them. The nanotubes were made from DLPC GUVs which are labeled by DiI-C20. The protocol is described in the method section. We did FCS measurements on both GUVs and nanotubes to compare the diffusion coefficients. As shown in Figure 8, the 2D FCS model fits to the correlation function of the GUVs very well. This fitting gives a diffusion coefficient of 1.06±0.01×10^-7^ cm^2^/s. By using this diffusing coefficient, we were able to predict the correlation function in one dimension (the dark green line in Figure 8). The experimental correlation function of the nanotubes is shown as circles in Figure 8. It falls between the 2D and 1D models, consistent with the result shown in Figure 5. Fitting this correlation function with our model gives a diffusion coefficient of 1.02±0.06×10^-7^ cm^2^/s, which is very similar to the values obtained from the GUVs. Furthermore, the fitting also gives a tube radius of 295±16 nm, which is consistent to our estimate from the fluorescence image (275-330 nm).

**Figure 8.**
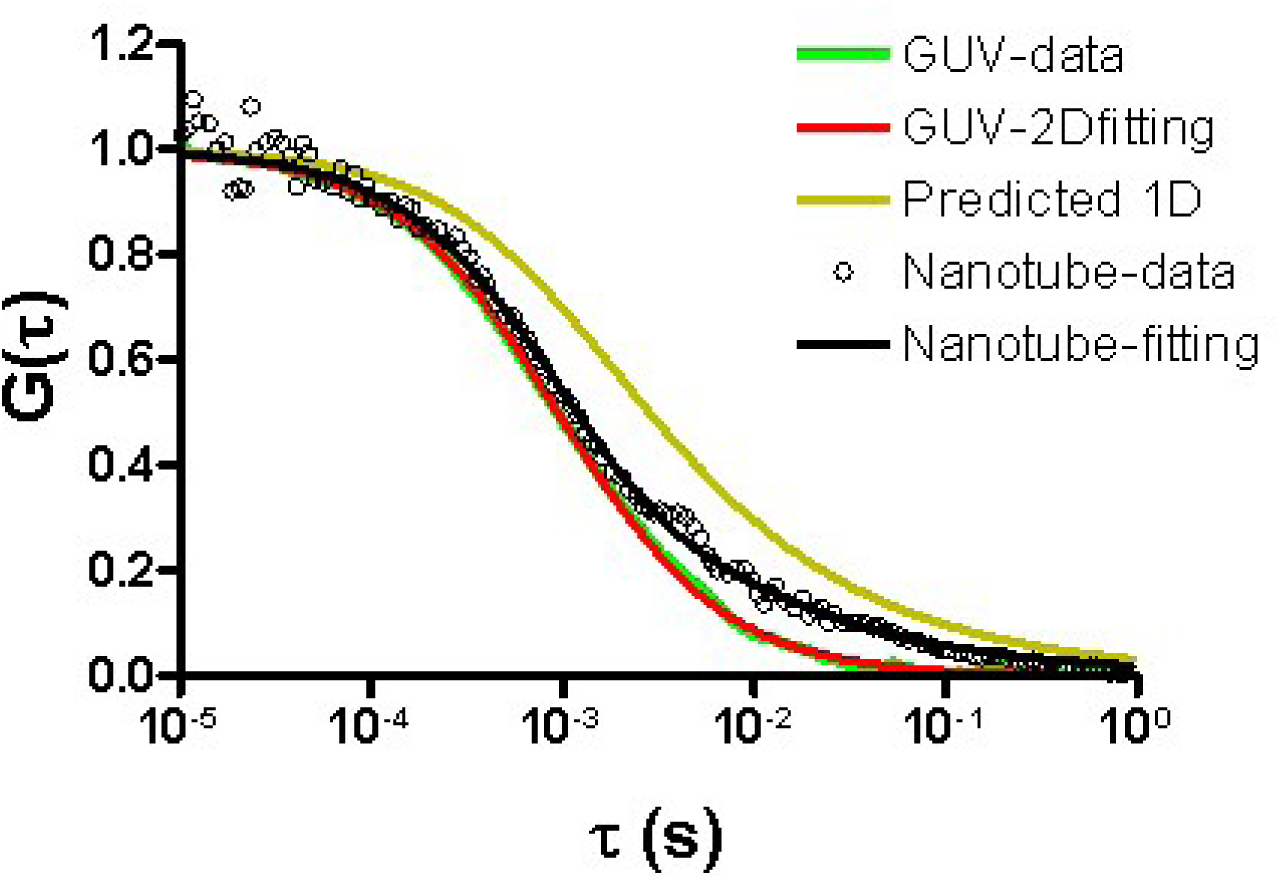
FCS for nanotubes comparing 2D and 1D diffusion. The bright green line is the experimental correlation function obtained from GUVs and the red line is the theoretical fitting with the 2D FCS model. The dark green line is the predicted 1D correlation curve based on the diffusion coefficient given by the 2D fitting (red). The circles define the experimental correlation curve obtained from nanotubes with the same composition as the GUVs described earlier (only DLPC). The black line is the fitting to the nanotube data with our model.

## 5. Discussion

### 5.1 Photon counting histogram in the closed system

In 1999, two different groups developed two similar techniques for analyzing the distribution of photon counts/intensity. One of them is called photon counting histogram (PCH)(Chen, Muller et al. 1999) and the other one is called fluorescence intensity distribution analysis (FIDA)(Kask, Palo et al. 1999). These two techniques are mathematically related (Meng and Ma 2006). However, there are still differences between them. One difference is that the generating function is used for FIDA. In other words, FIDA is the Fourier transform of PCH. Beside this difference, there is another important difference lying in the derivation procedures of these two methods. For PCH, the PCH for one molecule is first calculated. The fluctuation for one molecule is due to the unevenness of the illumination. Then the PCH for *n* molecules is the convolution of the PCH for one molecule by *n* times. In comparison to this, FIDA considers the fluctuation in a small volume (*dV*) first. The overall distribution is the convolution of the distributions for all *dV*s, leading to the sum of the logarithm of the generating function and therefore an integration over *V*. In the following context, I compare the two strategies for PCH and FIDA convolution of molecules and convolution of space, respectively. As we can see, PCH theory assumes that the distribution for different molecules are independent on each other, while the FIDA theory takes the assumption that the distribution for different positions are independent from other positions. Although these two procedures give the same result for open systems, FIDA is not suitable for closed systems. The reason is the distribution for different positions are not independent from each other for closed systems. For open systems, the distribution of the number of particles at on position is a Poisson no matter how many particles there are in other positions. However, in the closed system, the total number of particles is constant and all the positions compete with each other to have particles. Therefore, the convolution for all *dV*s is not suitable.

Chen et al (Chen, Muller et al. 1999), have already derived the PCH for closed systems. Here, we derived the Fourier transform of it. For one molecule in a closed system, the photon counting distribution is,

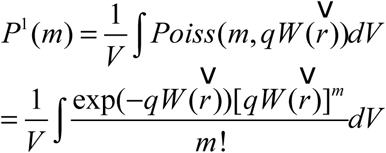

*m* is the number of photons.

Therefore, the generating function is

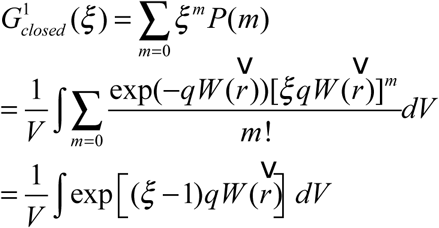

It can be easily shown that the distribution goes to a Poisson when 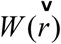 is constant over 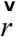, meaning the fluctuation is only due to the shot noise.

The generating function compares to the form for open systems (Kask, Palo et al. 1999):

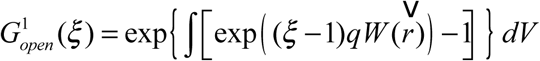

A fast way to calculate the generating function is to expand it into Talor series,

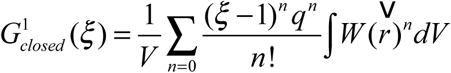

It compares to the logarithm of the generating function for open systems (Kask, Palo et al. 1999):

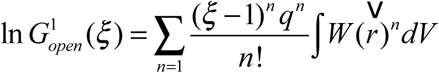

For *n* molecules

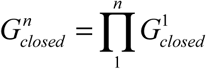

### 5.2 The problem of convolution of space

In retrospect to FCS, the problem of convolution of space also exists. This is the reason why the above correlation functions do not reach to zero at infinite time. To get *p*(*r*, *r'*,*τ*), the diffusion equation 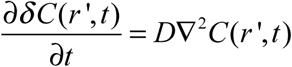 is solved with the initial condition *δ C*(*r*, 0)*δ C*(*r'*, 0) = *Cδ*(*r* – *r'*). (see Appendix). This initial condition means the concentration fluctuations for different positions are uncorrelated. However, this initial condition is only valid for the open system. Since for closed systems, different positions compete with each other to have molecules, there is a negative correlation between different positions. The correct initial condition should be 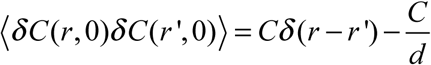. Therefore, equations can be revised as

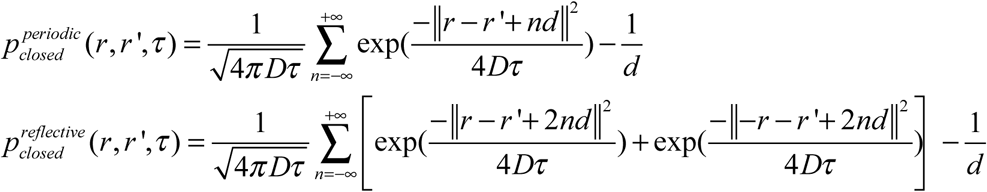

The derivation can also be performed using the convolution of molecules strategy as used for PCH. This approach will also give correlation functions that do not need to be forced to 0. See Appendix II.

### 5.3 Example for reflective boundaries

The theory for reflective boundaries can be useful for some single molecules studies. As depicted in Figure 9, we give two potential applications in the single molecule study.

**Figure 9.**
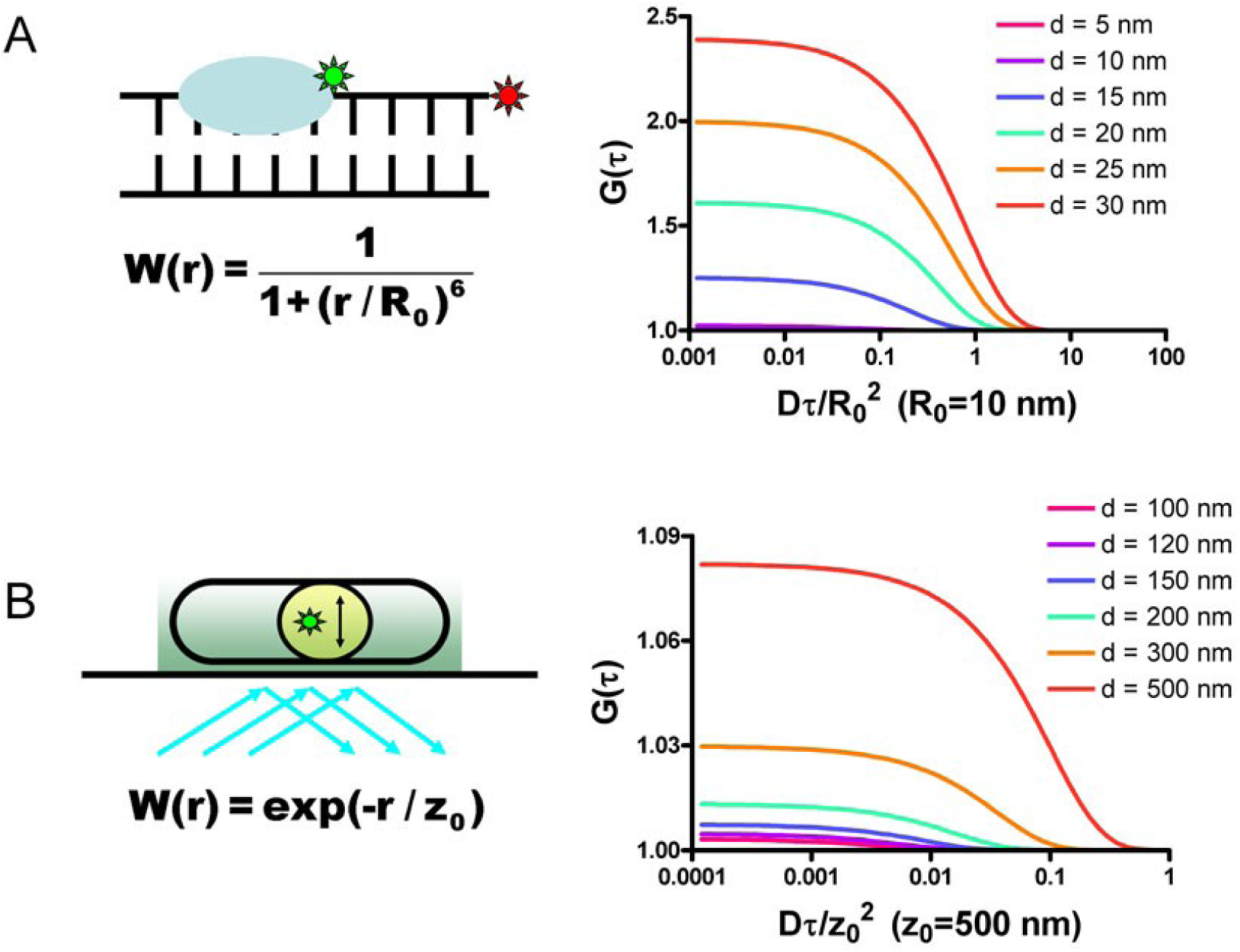
examples of single molecule experiments for which our theory could be useful. (A) protein diffusion along a short DNA. (B) diffusion in a confined region in the cell studied using TIRF (total internal reflection fluorescence microscopy).

The diffusion of proteins along DNA is usually studied with very long DNA molecules whose length is much longer than the optical resolution. To obtain the diffusion rates, a fitting step is required to get the precise position of the proteins. This is also the basic principle of the STORM and PALM super-resolution microscopy. One of the drawbacks is that the fitting step is time consuming. Here, we propose a single molecule experiment in which short DNA can be used (Figure 9A). For this experiment, a pair of FRET fluorphores are conjugated on the proteins and the end of the DNA, respectively. The diffusion of protein leads to a change of the distance between the FRET pair and causes a fluctuations of the detected intensity. The correlation function can be calculated and fitted with our theory for the reflective boundaries. In this case, the illumination function 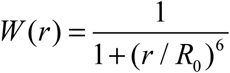, in which *R*_0_ is the Foster distance of the FRET pair.

Bacterial cells provide another popular field for single molecule studies. A typical E.coli cell spans 2 micron and has a 1 micron diameter. The diffusion of molecules in the cells is usually measured using TIRF (total internal reflection fluorescence microscope) in the x-y directions which are parallel to the surface. However, if the diffusion is confined in a region that is smaller than the optical resolution, it is difficult to obtain the diffusion rates. Here, we proposed a method to study the diffusion in the z-direction taking advantage of the decaying excitation intensity for TIRF (Mattheyses, Simon et al.). As shown in Figure 9B, as the protein moves in the z-direction, the brightness will change. The intensity can be recorded and the correlation function can be calculated and fitting with our theory. In this case, the illumination intensity decays exponentially. *W*(*r*) = exp(–*r* / *z*_0_)

As shown in Figure 9, for reflective boundaries, the apparent diffusion time (G(t) decay to half of the G(0)) increases with the size of the system, as for periodic boundaries. More prominently, the zero time amplitude change increases with the size of the system. This is because there is only one molecule for single molecule study.

## Appendix Appendix I: The correlation function calculated using in the Fourier space

*P*_*open*_ (*r*, *r'*,*τ*) is the solution of the diffusion equation 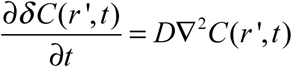 divided by C with the initial condition *δ C*(*r*, 0)*δ C*(*r'*, 0) = *Cδ*(*r* – *r'*). But since the expression is already known, *p*_*open*_ (*r*, *r'*,*τ*) can be directly calculated by finding the inverse Fourier transform of its Fourier transform. Therefore, the following expression is obtained.

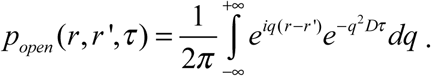

Substituting equation 1.1 into 10, the following equation can be obtained:

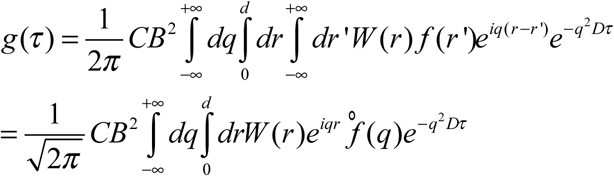

Because *f*(*r*) is periodic, its Fourier transform ° *f*(*q*) is composed of infinite series of pulses.

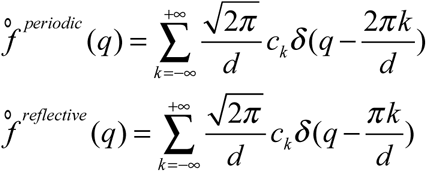

For the periodic boundaries, the factor 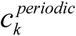 is

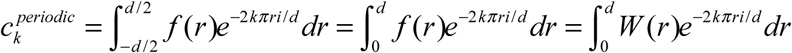

For reflective boundaries, *f*(*r*) can be expressed in cosine series only. Therefore,

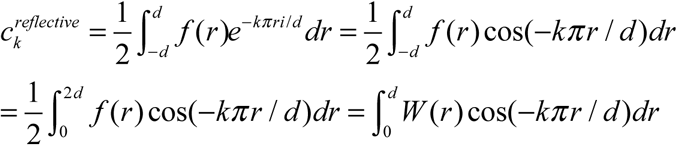

Therefore, for periodic boundary conditions,

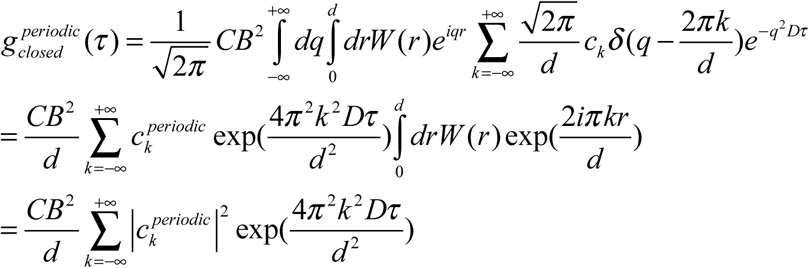

In the same way, the expression for the reflective boundaries can be obtained.

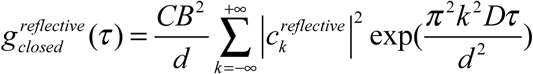

## Appendix II: proof of the initial condition 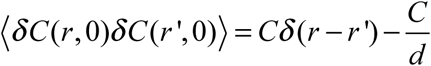

(1) First, we consider the case when *r* = *r*’, then 〈*δ C*(*r*, 0)*δ C*(*r*’, 0)〉 = 〈*δ C*(*r*, 0)^2^〉, which is the variance of the number of molecules at position *r*. For the closed system whose size is *d*, the distribution of the number of molecules at any positions should be a binomial distribution, 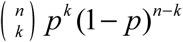, with *n* = *N*_*total*_ and 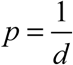. So the variance is

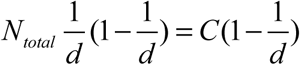

(2) Second, when *r* ≠ *r*’, a new random variable is introduced: the sum of the number of molecules at two positions *C*(*r*, *r*’, 0) = *C*(*r*, 0) + *C*(*r*’, 0). *C*(*r*, *r*’, 0) also follows the binomial distribution while the *n* = *N*_*total*_ and 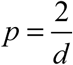. Therefore, the variance of *C*(*r*, *r* ’, 0) is 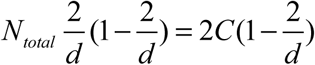. The variance of the sum of two random variables is:

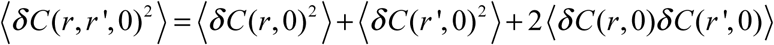

Therefore,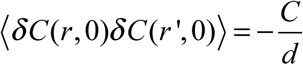

(3) Combind equations and. 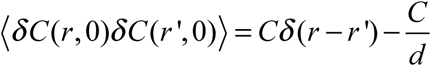

## Appendix III: calculate the correlation function through the convolution-of-molecules strategy

First, for any single molecule *i*, the probability that it is at position r is 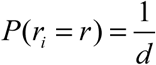, and the probability that it diffuses to position r’ after a lag time is then *p*_*closed*_ (*r*, *r* ’,*τ*) as indicated in equations (6) and (7). At any positions *r*, the fluorescence from single molecule is *BW*(*r*).

Therefore, the mean fluorescence of a single molecule is 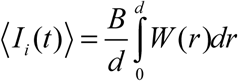, and 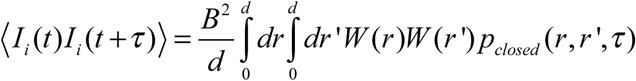, so

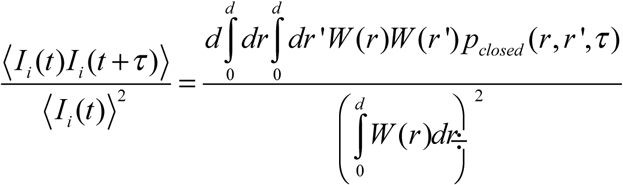

(2) Second, the total fluorescence intensity is usually calculated by *I*(*t*) = *B*∫*W*(*r*)*C*(*r*, *t*)*dr* for the convolution-of-molecules strategy. Here, instead of integrating over space, the intensity can also be calculated by summing up the contributions from all the molecules. 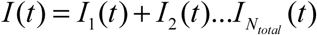,and *I*_*i*_ (*t*) = *BW*(*r*(*t*)) for each individual molecules. The fluctuation can be calculated in the same way: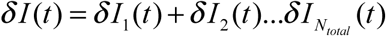. Here *δI*_*i*_ (*t*) = *δ I*_*i*_(*t*) – 〈*I*_*i*_〉. The mean in this equation is the mean fluorescence from individual molecules, therefore 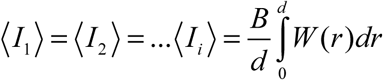.

For the correlation function 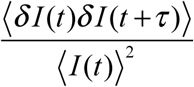, the denominator

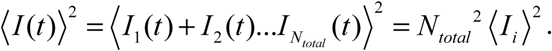

The nominator 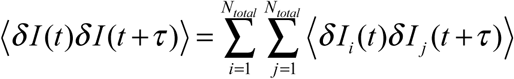

Because different molecules are uncorrelated,

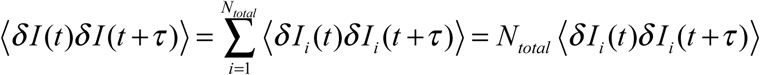

Combining equations and,

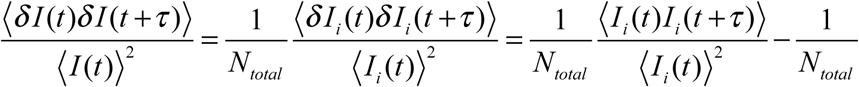

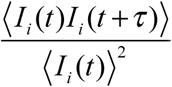 is given by equation.

It can be noticed in equation that the term 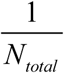 is already subtracted and the correlation function approaches to zero.

